# Characterization of Chronotypes Using the Symbolic Aggregate apprXimation (SAX) on Actigraphy Data

**DOI:** 10.1101/2024.09.03.611014

**Authors:** Wen Luo, Ioannis P. Androulakis

## Abstract

In this study, we discuss an efficient approach to characterizing chronotypes using Symbolic Aggregate approXimation (SAX) on actigraphy data. Actigraphy, a non-invasive monitoring of human rest/activity cycles, provides valuable insights into sleep-wake behaviors and circadian rhythms. However, the high dimensionality of actigraphy data poses significant challenges in storage, processing, and analysis. To address these challenges, we applied the SAX algorithm to transform continuous time-series actigraphy data into a symbolic representation, enabling dimensionality reduction while preserving essential patterns. We analyzed actigraphy data from the National Health and Nutrition Examination Survey (NHANES) database, covering over 10,000 individuals, and used unsupervised clustering to identify distinct chronotype patterns. The SAX transformation facilitated the application of machine learning techniques, revealing five chronotype clusters characterized by differences in activity onset, resolution, and intensity. Age distribution analysis showed biases towards specific age groups within the clusters, highlighting the relationship between age and chronotype. Key findings include age-related Chronotype variations with younger individuals exhibiting delayed chronotypes with significant differences in sleep onset (SOT) and wake time (WT) compared to older adults, suggesting a phase delay in sleep patterns as age decreases and activity transition dynamics where clusters showed distinct patterns in winding up and winding down periods, providing insights into the dynamics of activity transitions. This study demonstrates the efficiency and effectiveness of SAX in processing large-scale actigraphy data, enabling robust chronotype characterization that can inform personalized healthcare and public health initiatives. Further exploration of SAX integration with other biometric measures could deepen our understanding of human circadian biology and its impact on health and behavior.

## Introduction

Chronotype refers to an individual’s natural preference for sleep and wake times, often categorized into morningness and eveningness. It reflects the underlying circadian rhythm, which genetic, environmental, and behavioral factors influence. Understanding chronotypes is crucial for optimizing personal health, productivity, and well-being. Actigraphy, a non-invasive method of monitoring human rest/activity cycles, provides a valuable tool for inferring chronotypes. Actigraphy can offer insights into sleep-wake behaviors and circadian rhythms by capturing continuous, objective data on movement patterns. Actigraphy involves using a wrist-worn device that records movement over extended periods, typically resembling a watch. These devices contain accelerometers that detect and log physical activity levels, allowing researchers to infer periods of sleep and wakefulness. Actigraphy is widely used in sleep research due to its convenience, affordability, and ability to collect data over long durations in naturalistic settings (Fekedulegn et al., 2020). Unlike self-reported sleep diaries or questionnaires, actigraphy provides objective and continuous measurements, minimizing recall bias and offering a more accurate depiction of sleep patterns. Actigraphy can identify habitual sleep and wake times, critical markers of chronotype. Researchers can classify individuals into different chronotype categories by analyzing sleep periods’ timing, duration, and variability. Morning chronotypes typically show earlier sleep onset and wake times, while evening chronotypes have delayed sleep patterns. Beyond sleep-wake timing, actigraphy captures overall activity rhythms throughout the day. These rhythms, reflecting an individual’s energy levels and activity preferences, provide additional information on chronotype. For instance, morning chronotypes may exhibit higher activity levels in the early part of the day, while evening chronotypes might be more active in the evening. Actigraphy data also reveal sleep quality and regularity, which are influenced by chronotype. Evening chronotypes, for example, may experience more irregular sleep patterns and lower sleep quality due to social and work schedules misaligned with their natural preferences. Actigraphy can quantify these variations, offering a comprehensive view of how chronotype affects sleep health.

Actigraphy provides objective, quantifiable data, reducing the reliance on subjective self-reports, which can be prone to bias and inaccuracies. This objectivity enhances the reliability of chronotype assessments. The ability to continuously monitor over days, weeks, or even months allows for a thorough examination of an individual’s sleep-wake patterns and activity rhythms. This long-term data collection captures variations and trends that short-term studies might miss. Actigraphy allows data collection in real-world settings, reflecting typical daily routines and behaviors (Nikbakhtian et al., 2021). This ecological validity is crucial for understanding chronotypes in everyday life. Actigraphy-based chronotype identification is an excellent tool for large-scale studies where variability in chronotype is expected (Schneider et al., 2022)

Understanding an individual’s chronotype through actigraphy can inform personalized approaches to healthcare, such as optimizing timing for medication, therapy, and lifestyle interventions (Montaruli et al., 2021). Aligning work and school schedules with chronotype can enhance productivity, performance, and well-being (Su et al., 2022; Shim et al., 2023). For instance, flexible work hours or later school start times could benefit evening chronotypes. Population-level chronotype data can guide public health initiatives to improve sleep health and circadian alignment. Actigraphy-based research can identify at-risk groups and inform targeted interventions. Actigraphy is a powerful tool for inferring chronotypes, offering objective, continuous, and ecologically valid data on sleep-wake patterns and activity rhythms. By enhancing our understanding of chronotypes, actigraphy can contribute to personalized healthcare, optimized schedules, and improved public health outcomes. As technology advances, the integration of actigraphy with other biometric measures promises even deeper insights into human circadian biology and its impact on health and behavior.

The rise of wearable technology has revolutionized the collection of continuous physiological data, such as actigraphy, which records physical movement to monitor sleep-wake patterns and activity levels. Actigraphy generates vast data, often leading to storage, processing, and analysis challenges. Several computational approaches have been proposed for the statistical analysis of physical activity based on actigraphy (Krafty et al., 2019; Zhang et al., 2019), which is a subset of a larger problem focusing on the analysis of time series data (Bagnall et al., 2017). Dimensionality reduction through symbolic transformation offers a promising solution to these challenges, enabling more efficient and effective use of machine learning (ML) techniques. Actigraphy devices produce time-series data characterized by high dimensionality, with readings often recorded every minute or even more frequently. The sheer volume of data points can overwhelm traditional analytical methods and computational resources. High-dimensional data also poses challenges, such as the curse of dimensionality, where the performance of ML algorithms deteriorates due to the sparsity of data points in high-dimensional spaces. Symbolic transformation (Daw et al., 2003; Lin et al., 2003; Androulakis et al., 2007) involves converting continuous time-series data into discrete symbols, effectively reducing the dimensionality of the data while preserving essential patterns and structures. This process results in a symbolic representation that is more compact and manageable, offering opportunities for:

- Data Compression: Symbolic transformation significantly reduces the size of actigraphy data by converting continuous values into a sequence of symbols. This compression reduces storage requirements and speeds up data processing, making handling large datasets typical in wearable technology applications feasible.
- Noise Reduction: The symbolic representation can filter out minor fluctuations and noise inherent in raw actigraphy data, capturing the underlying trends and patterns more effectively. This denoising enhances the quality of the input data for ML models, leading to more robust and accurate predictions.
- Feature Extraction: Symbolic transformation facilitates extracting meaningful features from actigraphy data. Patterns, motifs, and anomalies become more apparent in the symbolic domain, enabling the identification of relevant features that can improve the performance of ML algorithms. For instance, repetitive patterns indicative of regular sleep cycles or activity bursts can be easily detected. Symbolic transformation mitigates the curse of dimensionality by reducing dimensionality, allowing ML algorithms to perform better. With fewer dimensions, algorithms can operate more efficiently, require less computational power, and exhibit improved accuracy and generalization. The symbolic representation provides a more interpretable format for understanding actigraphy data. Symbols corresponding to different activity levels or sleep states can be easily visualized and analyzed, aiding in interpreting model outputs and facilitating the communication of findings to non-experts.

Machine learning (ML) algorithms can leverage symbolic representations to classify daily activities like walking, running, or sedentary behavior. Dimensionality reduction using symbolic transformation of actigraphy data offers a powerful approach to overcoming the challenges posed by high-dimensional time-series data. Symbolic transformation enables more efficient and effective application of ML techniques by compressing, denoising, and extracting meaningful features. This synergy between symbolic transformation and ML holds great potential for advancing research and applications in health monitoring, sleep analysis, and beyond, making complex actigraphy data more manageable and actionable. Such applications would be invaluable in health monitoring, fitness tracking, rehabilitation, and, more significantly, chronotype inference. ML models can infer an individual’s chronotype by analyzing the symbolic patterns of activity and rest, providing insights into their circadian rhythms and optimizing schedules for better health and productivity.

In the present work, we analyzed actigraphy data from the National Health and Nutrition Examination Survey (NHANES) database, encompassing over 10,000 individuals, and applied unsupervised clustering to uncover distinct chronotype patterns. Applying the Symbolic Aggregate aproXimation (SAX) transformation facilitated machine learning techniques, identifying five chronotype clusters characterized by variations in activity onset, resolution, and intensity. An age distribution analysis revealed biases towards specific age groups within these clusters, underscoring the relationship between age and chronotype. Notably, younger clusters exhibited delayed chronotypes with significant differences in sleep onset (SOT) and wake time (WT) compared to older clusters, suggesting a phase delay in sleep patterns as age decreases. Additionally, the clusters displayed distinct patterns in winding up and winding down periods, providing valuable insights into the dynamics of activity transitions. This study demonstrates the efficiency and effectiveness of SAX in processing large-scale actigraphy data, enabling robust chronotype characterization that can inform personalized healthcare and public health initiatives.

## Methods

### Data

We extracted physical activity monitoring data from the NHANES database. Specifically, we considered recordings from the 2011^1^ and 2013^2^ surveys. The actigraphy data is provided per minute (variable name: PAXMTSM). The nationwide, cross-sectional surveys contained recordings for 6,710 (2011) and 7,401 (2013) individuals, respectively. Thus, our study considered 14,111 recordings of actigraphy data over seven days. Not all individuals had complete records. However, our approach considers an average day, so missing data were averaged. The collection process details are standardized and have already been described in earlier publications(Su et al., 2022; Shim et al., 2023) and will not be repeated here.

### Transformation and symbolic representation

In this work, we propose using the Symbolic Aggregate aproXimation (SAX) algorithm (Lin et al., 2003; Lin et al., 2007). SAX is a technique for transforming time series data into a symbolic representation, enabling efficient data mining and analysis. The process involves several key steps:

a. Normalization: The time series data is normalized to have a mean of zero and a standard deviation of one. This step ensures the data is on a standard scale, facilitating meaningful comparisons.
b. Segmentation: The normalized time series is divided into equal-length segments. Each segment represents a portion of the time series data.
c. Dimensionality Reduction: Each segment is transformed into a single value using Piecewise Aggregate Approximation (PAA) techniques. PAA reduces the dimensionality of the data by averaging the values within each segment, resulting in a lower-dimensional representation of the original time series.
d. Symbolic Mapping: The reduced-dimensional representation is then converted into a symbolic representation. This is achieved by mapping the segment averages to discrete symbols based on predefined breakpoints. These breakpoints are determined using the normal distribution to ensure equal-probability segments.
e. String Formation: The symbols corresponding to each segment are concatenated to form a symbolic string. This string is a compressed and discrete representation of the original time series, capturing its essential patterns and trends.

The SAX algorithm is beneficial for long time series analysis because it allows for efficient indexing, comparison, and clustering of time series data. The symbolic representation makes applying string-based algorithms to time series data possible, facilitating motif discovery, anomaly detection, and pattern-matching tasks. SAX’s ability to reduce dimensionality and provide a symbolic representation while preserving the critical characteristics of the original time series makes it a powerful tool in data mining and analysis. We have previously applied the method to analyze genome-wide longitudinal transcriptomic data (Yang et al., 2008; Scheff et al., 2010).

The methodology is succinctly presented in **Figure 1**. According to the study design, activity data is recorded at 1 min resolution, and the wearable devices are used for a maximum of seven days. The NHANES Survey objectively measures physical activity using a Physical Activity Monitor (PAM). This device recorded acceleration across three axes (x, y, and z) at a frequency of 80 Hz, along with ambient light levels at 1 Hz. In the NHANES dataset, PAXMTSM represents each minute’s MIMS (Motion Intensity Measurement Summary) triaxial value. It is a summary measure derived from accelerometer data collected by physical activity monitors (PAMs) worn by participants. The PAXMTSM variable is calculated by summing the minute summary acceleration measurements obtained on the x-, y-, and z-axes (PAXMXM, PAXMYM, and PAXMZM). It represents the total movement intensity recorded during that minute.

**Figure 1:**
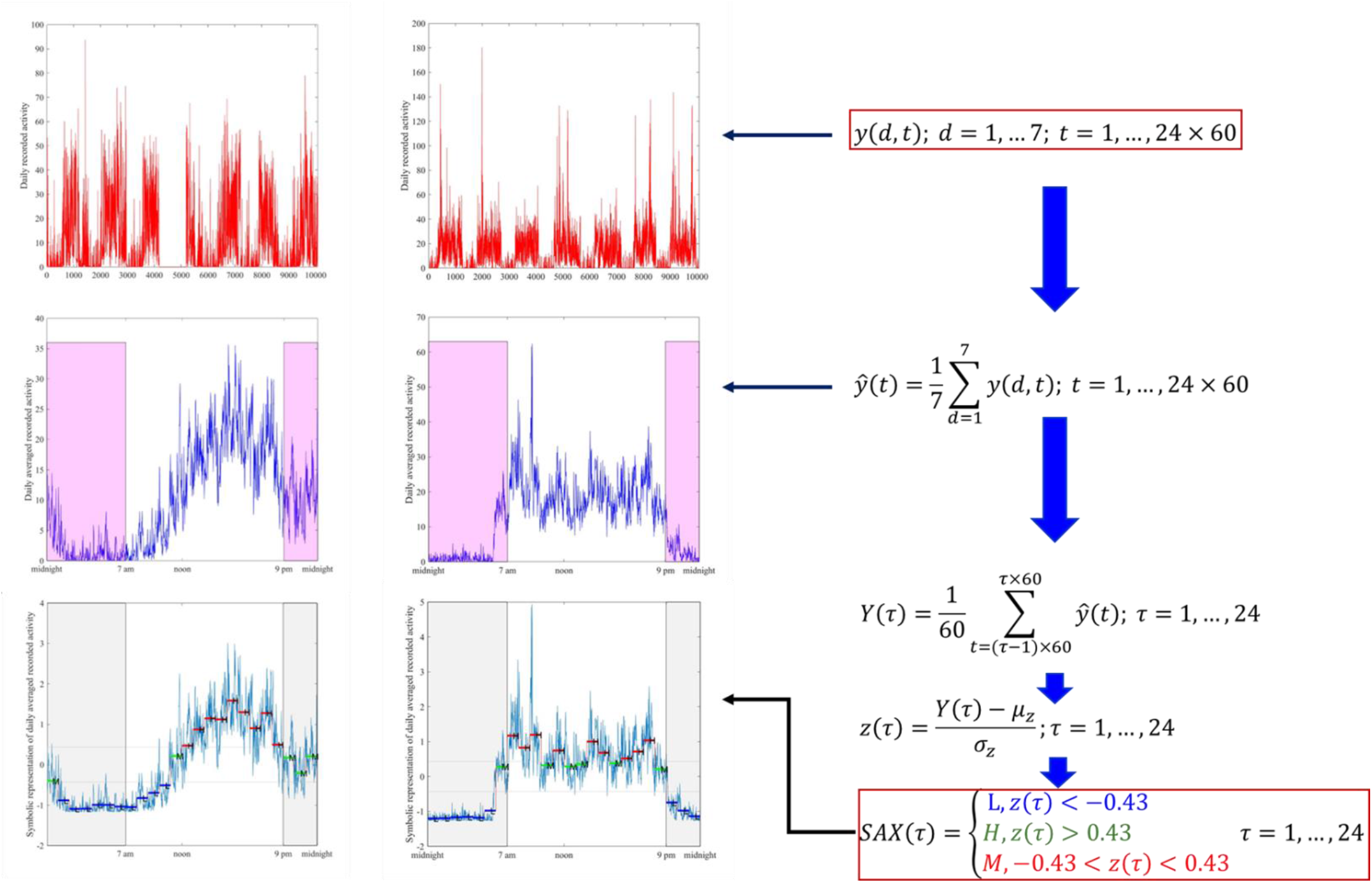
NHANES actigraphy data records, at most, seven days of activity data at a resolution of 1 minute. The maximum number of recorded values *y*(*d, t*), *d* = 1, …, 7; *t* = 1, …, 24× 60 per subject would be 7× 24× 60. The top panels depict two such recordings for two subjects. The multiple-day recordings are averaged per minute to form an “average day,” as shown in the middle panels. Finally, the average day is processed according to SAX by first averaging data over 1-hour intervals, thus reducing the effective dimensionality to 24. The data is subsequently z-scored and then converted to a symbolic representation. Finally, the original data set is represented by 24 symbols.

As shown in **Figure 1**, the raw data (top panels) record activity over several days. For illustration purposes, we show recordings of two different subjects. Usually, recordings are reported over seven days. However, since we will eventually describe an average day, the number of days is unimportant. The recordings are reported at the rate of one per minute. Therefore, there are 24× 60 = 1440 daily recordings, one for each minute of the 24-hour day. Subsequently (middle panels), the recording at each minute of the day is averaged over the number of days for which recordings exist. We do so because we treat each day as a replicate. In doing so, we may lose day-to-day variations. However, because of the short timespan for recordings, it is hard to differentiate between errors and persistent patterns (such as shift work). The method would still be valid without losing generality if one considered daily variations (this issue will be further discussed later.)

Once the data is averaged over the available days, an “average” day’s actogram is developed. At this point, the steps of SAX can be applied: each day is normalized to have zero mean and 0 standard deviation; the data is segmented such that activity is reflected per hour. This segmentation aims to smooth the data and eliminate low-level noise. Once the data has been segmented, the symbolic transformation is applied, which converts each time series to a sequence of symbols. In our analysis, we have used three levels: Low, Medium, and High. Experimentation indicated that this level of approximation produced consistent results. The breakpoints (*β*_1_, *β*_2_) are determined such that

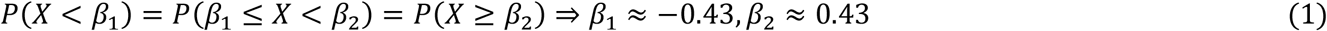

The steps are summarized in the bottom part of **Figure 1.** The power of SAX is that it allows for a substantial dimensionality reduction while maintaining the critical characteristics of the original dynamics. To infer chronotype characteristics, the approach retains all the critical properties at an appropriate level of granularity. The 1-hr segmentation effectively filters, reducing the noise associated with the measurements. A key advantage of SAX is its ease of implementation.

Once all of the subjects have been transformed into their symbolic representation, many machine learning algorithms can be applied to all kinds of analyses. In the present work, we primarily examined whether specific activity patterns emerge from the recorded data. Therefore, we were interested principally in unsupervised clustering of actograms. In this study, we utilized the elbow method to determine the optimal number of clusters for analyzing our actigraphy data. The elbow method is a heuristic approach in cluster analysis beneficial for k-means clustering, where the optimal number of clusters must be specified a priori. We defined a range of possible cluster counts (k) from 1 to 10. For each k, we applied the k-means algorithm to our dataset and computed the within-cluster sum of squares (WCSS), quantifying each cluster’s variance. The WCSS was recorded for each k, resulting in a vector of WCSS values. To identify the optimal number of clusters, we calculated the first and second derivatives of the WCSS vector. The first derivative represents the change in WCSS as k increases.

In contrast, the second derivative indicates the rate of change of this slope, effectively capturing the curvature of the WCSS plot. The point of maximum curvature, or the “elbow” point, typically signifies a significant reduction in the rate of decrease of WCSS, suggesting a balance between the goodness of fit and the complexity of the model. We identified this elbow point by locating the k value before the sign of the second derivative changes, corresponding to the second derivative’s minimum absolute value. This k value was adjusted to account for the length reduction caused by the differencing operations. We plotted the WCSS against the number of clusters to validate our findings, visually confirming the elbow point. Subsequently, we performed the final k-means clustering using the determined optimal number of clusters. This methodological approach ensures that the clustering model is both parsimonious and effective, capturing the intrinsic structure of the actigraphy data while avoiding overfitting. The elbow method, therefore, provided a robust framework for determining the appropriate number of clusters, crucial for accurately inferring chronotypes from the actigraphy measurements.

## Results

Combining the 2011-2013 data created a cohort of 10,016 individuals for which actigraphy and age data were available. The data represent a 50/50 split between males and females, ranging in age from 12 to 80, with a median age of 41. The clustering of the SAX transformed data is illustrated in the top part of **Figure 2**. The clustering identified five distinct chronotypes characterized by differences in the onset and resolution of activity and terms of the intensity of the activity. We subsequently mapped the original actigraphy data (daily averaged) onto the identified clusters as an implicit test. Interestingly, the clear separation is maintained, as seen in the bottom part of **Figure 2.** It is important to emphasize that this is not the result of “reclustering” the data but rather mapping the raw data onto the clusters identified by clustering the SAX transformed actigraphy data.

**Figure 2:**
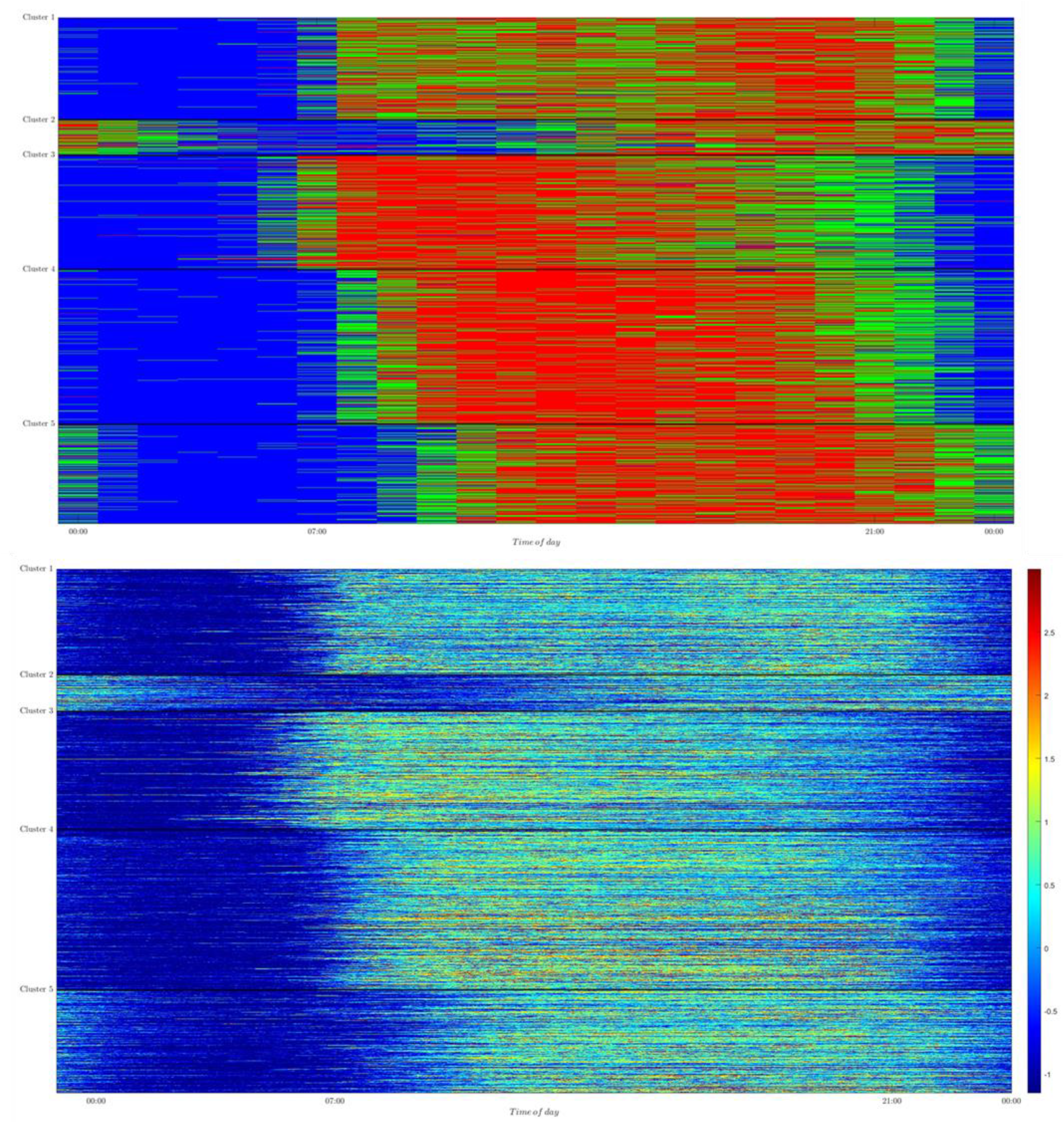
Clustering of SAX-transformed 10,016 actograms from the 2011-2013 NAHNES surveys. (top) Clustering of the SAX transformed acrograpms. Colors indicate high (red), middle (green), and low (blue) values; (bottom) Raw actigraphy data mapped onto SAX clusters. For the bottom graph, the actigraphy data were not reclustered; instead, the raw z-scored data were grouped according to the cluster assignment resulting from the SAX-transformed data. The data illustrate the method’s effectiveness since the top part clusters the 24 symbols per subject (row), whereas the bottom visualizes 1,440 points per subject (row).

To obtain a better sense and likely quantification of the sleep onset (SOT) and wake time (WT) as well as activity throughout the day characteristics of each cluster, we determined the SAX representation of the center of each cluster, **Figure 3**. For furthering the analysis, the waveform of each cluster center was approximated by a piecewise linear function:

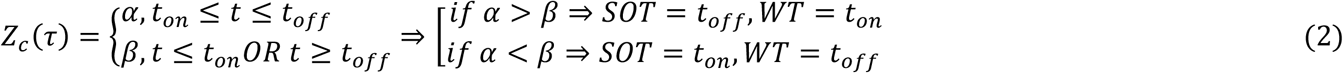

**Figure 3:**
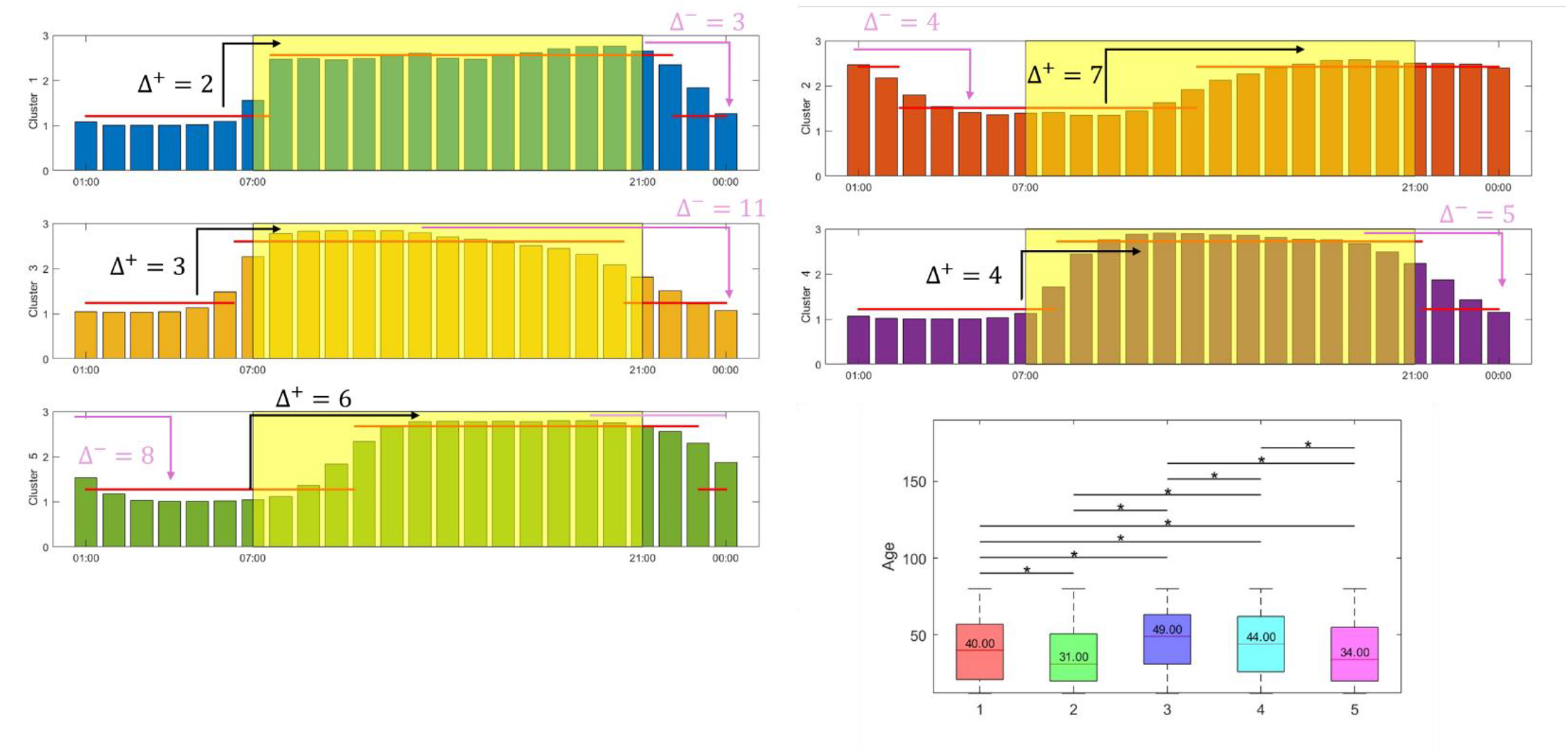
SAX-transformed activity levels of cluster centers. This representation enables a more nuanced characterization of the expected chronotype for each cluster. Interestingly, the late chronotypes (clusters 2 and 5) contained overwhelmingly younger (including adolescent) subjects. The distribution of the ages graph also indicates statistically significant (*P*_*val*_ < 0.05) age differences between groups. Interestingly, groups 2 and 5 (younger subjects)

This simple transformation allows us to gain a better insight into the characteristics of each cluster and, by extension, the members (subjects) belonging to it. The results are depicted in **Figure 3**, which denotes the hourly activity level for each cluster and the piecewise linear approximation of each cluster’s center. The characteristics of each cluster are shown in **Table 1**, which also depicts the median age of the subjects assigned to each group. The age distribution and differentiation of clusters based on these distributions are also shown in **Figure 3**.

**Table 1:** Cluster-specific chronotype attributes.

| Cluster | Median age | SOT | WT | $\alpha$ | $\beta$ | $t_{active}$ | $\Delta^+$ | $\Delta^-$ |
| --- | --- | --- | --- | --- | --- | --- | --- | --- |
| 1 | 40.0 | 22.11 | 7.63 | 1.21 | 2.56 | 14.48 | 2 | 3 |
| 2 | 31.0 | 2.46 | 13.15 | 2.42 | 1.51 | 13.31 | 7 | 4 |
| 3 | 49.0 | 20.34 | 6.34 | 1.23 | 2.60 | 14.00 | 3 | 11 |
| 4 | 44.0 | 21.24 | 8.14 | 1.22 | 2.72 | 13.10 | 4 | 5 |
| 5 | 34.0 | 23.03 | 10.68 | 1.27 | 2.67 | 12.35 | 6 | 8 |

## Discussion

In this study, we explored SAX as a means to reduce, in a meaningful way, the dimensionality of actigraphy data and demonstrate how these could be used to enable further machine learning tasks as well as generate meaningful insights into subjects’ behavior, with emphasis on chronotype. We examined publicly available data through the NHANES repository without loss of generality, focusing on two years’ worth of data representing over 10,000 individuals for which actigraphy data was available for, on average, seven days. Therefore, the dataset is not a trivial one.

We processed the data to establish a “representative” day of an individual’s activity by identifying the average activity levels over the period for which actigraphy data were available. We assume that each “day” is a “replicate” in a massive temporal experiment. There is no doubt that individuals may exhibit activity patterns from day to day. However, the seven-day period available through the NHANES data is not readily amenable to such an analysis. However, it must be emphasized that the study as presented can be trivially extended to account for each day separately. However, as mentioned, given the nature of the available data, that would probably not be the optimal approach. Nevertheless, the analysis can be performed but not discussed here so as not to distract from the key message of this work.

The symbolic transformation process is straightforward, which makes it generalizable and easily implementable. We chose an alphabet of three: low, mid, and high levels of activity, reflecting an appropriate balance between indicative or rest, activity, and transition. The achieved informative dimensionality reduction is substantial. Given that we consider segmentation of the data at 1-hour intervals, the reduction is a factor of 60 per day (assuming that the data is reported on a per-minute basis, as is in NHANES). The 1-hour interval was selected in our study as a reasonable way to differentiate between actual change in activity and “noise.” However, this is a variable that is easily adapted. However, for practical purposes, anything less than that would not introduce any meaningful information, given that overall activity changes at a scale of less than one hour would correspond to special cases. When dealing with peoples’ long-term habits (such as chronotype), it is essential to capture high-level behavior patterns.

The clustering results of **Figure 2** and the associated analysis presented in **Figure 3** are very informative. The symbolically transformed, daily averaged actigraphy profiles were clustered into five distinct groups. The optimal number of clusters was determined by applying the elbow method. The top part of **Figure 2** illustrates the partitioning of subjects based on group membership. The partitioning is validated in the bottom part, which depicts the grouping of the raw data when subjects are assigned to clusters as determined by the analysis of the symbolic transformations.

All groups exhibited an expected behavior at the highest level: relatively extended periods of high and low activity, separated by transition periods reflecting transitions from rest to active and vice versa. However, a closer examination of the results indicates some exciting patterns.

a. There is a clear bias towards age distribution in the actigraphy clusters, as noted in earlier studies (Neikrug et al., 2020). The age distribution across the clusters is compared using standard Anova and Tuckey-Kramer multiple comparison methods, and it was determined that the age distributions were different between all groups, except for clusters 2 and 5, i.e., the younger groups, **Figure 3**. This implies that the actigraphy data have dynamics biased towards age groups.
b. Clusters 2 and 5, which are populated primarily by adolescents and younger adults, as indicated by the median age (31 and 34, respectively), tend to exhibit delayed chronotypes with SOT 02:46 and 23:03, respectively, as approximated by equation (2), **Table 1**. Interestingly, the general trend is towards late chronotypes, which advance as the median age gets younger, **Figure 4**. It is interesting to see that the “younger” groups are broken down into two sub-groups: cluster 2, a true “night owl” with a SOT in the early morning hours (median age = 31), and a reasonable phase delayed group, cluster 5, with a SOT around 11 AM (median age = 34). Furthermore, the older group, cluster 3, had the earliest expected SOT, then SOT dropped almost linearly, **Figure 4**. A similar trend exists for the wake time, with WT being phase-delayed as the median age decreases.
c. Using the cluster center representations, we aimed to gain more insight into the details of the cluster-dependent activities. An interesting observation relates to the time required for a subject to reach peak activity levels, which we define as “winding up,” Δ^+^, and the “winding down” period, which we define as the time it takes to enter the rest period, Δ^−^. Based on the information in **Figure 3**Figure 3, we determined these quantities as shown in **Table 1**. A few interesting observations are in order:
  - Cluster 3 (median age 49) exhibits a relatively short winding-up time, Δ^+^ ≈ 3, but a protracted winding down, Δ^−^ ≈ 11.
  - Cluster 4 (median age 44) exhibits somewhat similar Δ^+^/Δ^−^ times of 4 and 5, they are respectively, implying more rapid relaxation than Cluster 3.
  - Cluster 1 (median age 40) appears to exhibit the more “conventional” activity patterns, with a WT around 08:00 and SOT around 20:00 appearing to exhibit relatively short and somewhat similar Δ^+^ and Δ^−^ values. Also, subjects in this cluster appear to maintain a relatively robust activity during the interval [WT, SOT]. Compared to clusters 3 and 4, the subjects in this cluster appear to be active for longer but also exhibit faster transitions (rest ↔active).
  - Cluster 5 (median age 34) is one of the two “younger” clusters, exhibiting similar Δ^+^/Δ^−^ times and relatively constant active periods. Finally
  - Cluster 2 (median age 31) is the “younger” cluster, which, interestingly, exhibits relatively protracted winding up and winding down periods.

**Figure 4:**
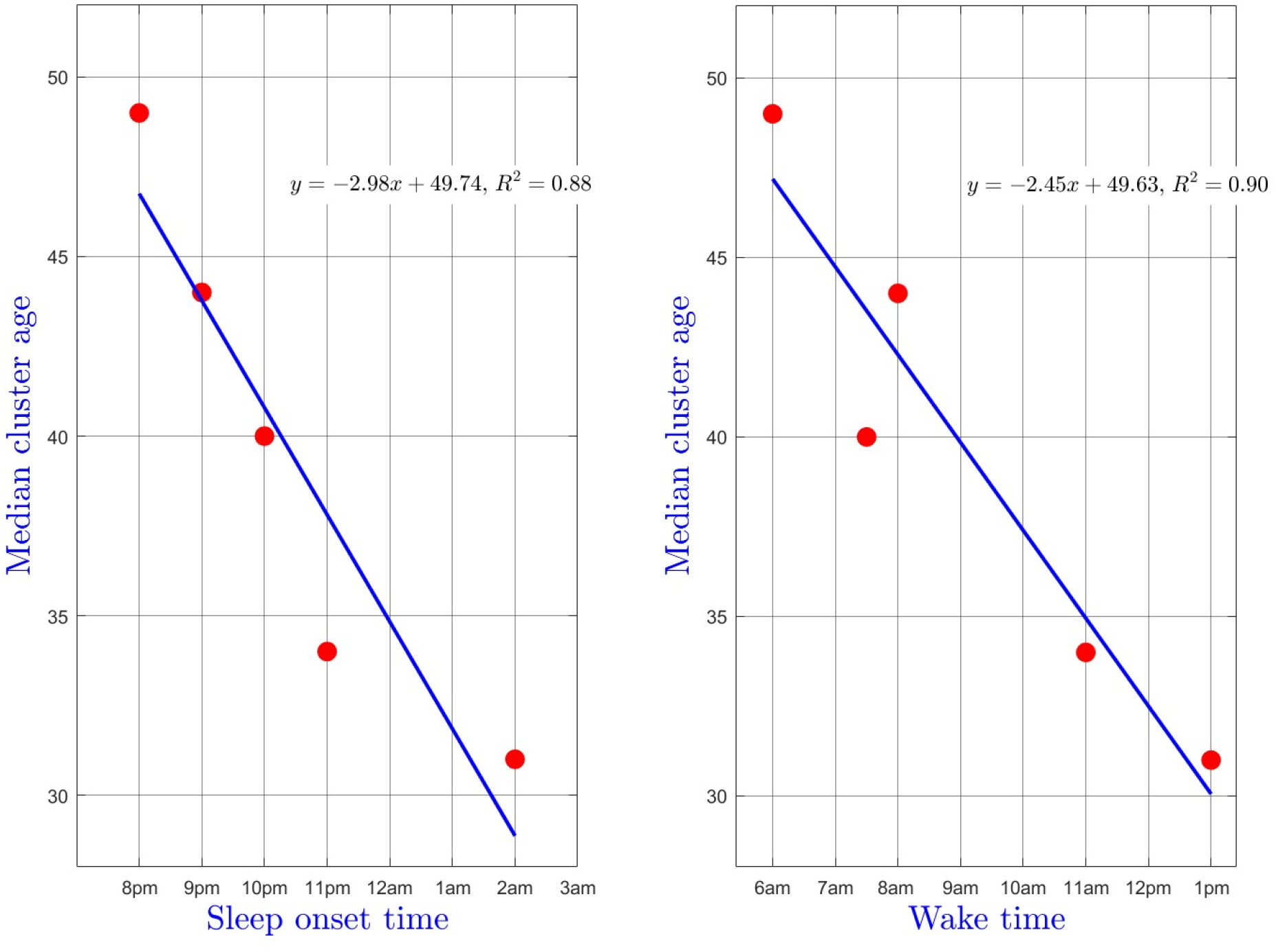
Cluster analysis revealed a noticeable phase delay in SOT and WT with age.

## Conclusions

In this study, we explored the use of Symbolic Aggregate approXimation (SAX) to efficiently reduce the dimensionality of actigraphy data and enhance the characterization of chronotypes. By analyzing data from the NHANES database, covering over 10,000 individuals, we demonstrated how SAX could transform high-dimensional time series data into a manageable and interpretable format. This transformation facilitated the application of machine learning techniques, particularly unsupervised clustering, to uncover distinct chronotype patterns in the population. Our methodology involved: Data Normalization and Symbolic Transformation, Clustering and Chronotype Characterization, and Cluster Analysis and Insights. Ourexamination of cluster centers provided insights into the activity patterns, sleep onset (SOT), wake time (WT), and transition periods. Age distribution analysis revealed biases towards certain age groups within the clusters, highlighting the relationship between age and chronotype. Our key findings point towards 1) age related chronotype variations with younger clusters (clusters 2 and 5) exhibiting delayed chronotypes, with significant differences in SOT and WT compared to older clusters. This trend suggests a phase delay in sleep patterns as age decreases; 2) Activity transition Dynamics where clusters showed distinct patterns in winding up and winding down periods. Overall the SAX-transformed data provided a robust framework for efficiently analyzing large-scale actigraphy datasets, enabling the identification of chronotype patterns that can inform personalized healthcare and public health initiatives. While our study highlights the efficiency and effectiveness of SAX in processing actigraphy data, several limitations should be noted: The NHANES dataset tends to overrepresent relatively healthy individuals, potentially skewing the results. This bias arises from voluntary participation, exclusion criteria, and self-selection, which may not accurately reflect populations with higher disease prevalence. Second, the methodology can be extended to account for daily variations, but our analysis focused on average activity patterns due to the seven-day recording period in NHANES data. Fnally, although SAX captures subtle nuances in activity patterns, further research is needed to establish the direct health implications of these variations.

In conclusion, this study demonstrates a powerful approach for efficiently analyzing actigraphy data to characterize chronotypes. The SAX method offers substantial dimensionality reduction while maintaining critical characteristics of the original time series, enabling meaningful insights into activity patterns and their relationship with chronotypes. Future work should explore the integration of SAX with other biometric measures to deepen our understanding of human circadian biology and its impact on health and behavior.

## Acknowledgment

IPA acknowledges support from MIH GM131800

## Footnotes

1 https://wwwn.cdc.gov/nchs/nhanes/search/datapage.aspx?Component=Examination&CycleBeginYear=2011

2 https://wwwn.cdc.gov/nchs/nhanes/search/datapage.aspx?Component=Examination&CycleBeginYear=2013

## References

Androulakis IP, Yang E, and Almon RR (2007) Analysis of time-series gene expression data: methods, challenges, and opportunities. Annu Rev Biomed Eng 9:205–228.

Bagnall A, Lines J, Bostrom A, Large J, and Keogh E (2017) The great time series classification bake off: a review and experimental evaluation of recent algorithmic advances. Data Min Knowl Discov 31:606–660.

Daw CS, Finney CEA, and Tracy ER (2003) A review of symbolic analysis of experimental data. Review of Scientific Instruments 74:915–930.

Fekedulegn D, Andrew ME, Shi M, Violanti JM, Knox S, and Innes KE (2020) Actigraphy-Based Assessment of Sleep Parameters. Ann Work Expo Health 64:350–367.

Krafty RT, Fu H, Graves JL, Bruce SA, Hall MH, and Smagula SF (2019) Measuring Variability in Rest-Activity Rhythms from Actigraphy with Application to Characterizing Symptoms of Depression. Statistics in Biosciences 11:314–333.

Lin J, Keogh E, Lonardi S, and Chiu B (2003) A symbolic representation of time series, with implications for streaming algorithms. In Proceedings of the 8th ACM SIGMOD workshop on Research issues in data mining and knowledge discovery, pp 2–11, Association for Computing Machinery, San Diego, California.

Lin J, Keogh E, Wei L, and Lonardi S (2007) Experiencing SAX: a novel symbolic representation of time series. Data Mining and Knowledge Discovery 15:107–144.

Montaruli A, Castelli L, Mule A, Scurati R, Esposito F, Galasso L, and Roveda E (2021) Biological Rhythm and Chronotype: New Perspectives in Health. Biomolecules 11.

Neikrug AB, Chen IY, Palmer JR, McCurry SM, Von Korff M, Perlis M, and Vitiello MV (2020) Characterizing Behavioral Activity Rhythms in Older Adults Using Actigraphy. Sensors (Basel) 20.

Nikbakhtian S, Reed AB, Obika BD, Morelli D, Cunningham AC, Aral M, and Plans D (2021) Accelerometer-derived sleep onset timing and cardiovascular disease incidence: a UK Biobank cohort study. Eur Heart J Digit Health 2:658–666.

Scheff JD, Almon RR, DuBois DC, Jusko WJ, and Androulakis IP (2010) A new symbolic representation for the identification of informative genes in replicated microarray experiments. OMICS 14:239–248.

Schneider J, Fárková E, and Bakštein E (2022) Human chronotype: Comparison of questionnaires and wrist-worn actigraphy. Chronobiology International 39:205–220.

Shim J, Fleisch E, and Barata F (2023) Wearable-based accelerometer activity profile as digital biomarker of inflammation, biological age, and mortality using hierarchical clustering analysis in NHANES 2011-2014. Sci Rep 13:9326.

Su S, Li X, Xu Y, McCall WV, and Wang X (2022) Epidemiology of accelerometer-based sleep parameters in US school-aged children and adults: NHANES 2011-2014. Sci Rep 12:7680.

Yang E, Maguire TJ, Yarmush ML, Berthiaume F, and Androulakis IP (2008) Identification of regulatory mechanisms of the hepatic response to thermal injury. Computers & Chemical Engineering 32:356–369.

Zhang Y, Li H, Keadle SK, Matthews CE, and Carroll RJ (2019) A Review of Statistical Analyses on Physical Activity Data Collected from Accelerometers. Stat Biosci 11:465–476.

